# Whole-chromosome fusions in the karyotype evolution of *Sceloporus* (Iguania, Reptilia) are more intense in sex chromosomes than autosomes

**DOI:** 10.1101/2020.03.31.011619

**Authors:** Artem P. Lisachov, Katerina V. Tishakova, Svetlana A. Romanenko, Anna S. Molodtseva, Dmitry Yu. Prokopov, Jorge C. Pereira, Malcolm A. Ferguson-Smith, Pavel M. Borodin, Vladimir A. Trifonov

## Abstract

There is a growing body of evidence that the common ancestor of vertebrates had a bimodal karyotype, i.e. consisting of large macrochromosomes and small microchromosomes. This type of karyotype organization is preserved in most reptiles. However, certain species independently experience microchromosome fusions. The evolutionary forces behind this are unclear. We investigated the karyotype of the green spiny lizard, *Sceloporus malachiticus,* an iguana species which has 2n=22, whereas the ancestral karyotype of iguanas had 2n=36. We obtained and sequenced flow-sorted chromosome-specific DNA samples and found that most of the microchromosome fusions in this species involved sex chromosomes. We found that certain ancestral squamate chromosomes, such as the homologue of the *Anolis carolinensis* chromosome 11, are repeatedly involved in sex chromosome formation in different species. To test the hypothesis that the karyotypic shift could be associated with changes in recombination patterns, and to study sex chromosome synapsis and recombination in meiosis, we performed synaptonemal complex analysis in this species and in *S. variabilis,* a related species with 2n=34. We found that in the species studied the recombination patterns correlate more with phylogeny than with the structure of the karyotype. The sex chromosomes had two distal pseudoautosomal regions and a medial differentiated region.

## Introduction

There are two main types of karyotype organization in vertebrates: unimodal and bimodal [1]. In bimodal karyotypes there is a clear distinction between normal-sized macrochromosomes and small “dot-like” microchromosomes. This is a characteristic feature for some amphibians and fishes (mostly basal clades), birds, and a majority of reptiles [2]. Unimodal karyotypes have chromosomes of gradually decreasing size. They are characteristic for many amphibians (derived clades), teleost fishes, mammals, and some reptiles [3]. The bimodal karyotype organization is thought to be ancestral for vertebrates, since the microchromosomes of different lineages share high homology [4,5]. The unimodal organization originated independently in different lineages by fusion of microchromosomes with each other and with the macrochromosomes [6,7].

This parallel process is interesting in the context of the search for general patterns of karyotype and genome evolution in animals. Fixation of chromosomal rearrangements is often presumed to be neutral and random, but the repetition of similar rearrangements points to their possible biological significance, as suggested by Morescalchi [1,8]. According to this hypothesis, decreasing the overall recombination rate via lowering the chromosome number under a selection for decreased recombination could be a possible biological mechanism for shifts to unimodality. In this case, it could be expected that other forces, such as changes in crossover rates, would also act to decrease recombination, and the species with unimodal karyotypes would have lower crossing over rates than related species with bimodal karyotypes. The molecular cytogenetic and genomic methods offer new possibilities to study the parallel shifts to unimodality.

To address the question of possible adaptive significance of microchromosome fusions, it is important to determine whether different microchromosomes fuse in random combinations, or if certain chromosomes tend to fuse with each other repeatedly in different lineages. This could happen if the linkage between the alleles of certain loci is favored by selection[1,9]. The fusions involving sex chromosomes are particularly interesting, since it has been suggested that certain chromosome pairs are more often involved in sex chromosome formation in the vertebrate evolution than others, both by becoming sex chromosomes *de novo* and by fusing with already existing sex chromosomes [10,11]. Mammals and teleost fishes experienced loss of microchromosomes early in their evolution, and subsequent chromosomal and genomic rearrangements obscured the initial steps of the microchromosome fusion [5]. Squamate reptiles, on the contrary, contain species with both bimodal and unimodal karyotypes, and some clades with unimodal karyotypes have formed recently and have close relatives with bimodal karyotypes.

The genus *Sceloporus* (Iguania, Pleurodonta, Phrynosomatidae) displays higher chromosomal variability than reptiles on average: 2n varies in this genus from 22 to 46. All these karyotypes can be derived from the ancestral phrynosomatid karyotype (2n=34) via chromosomal fusions and fissions[12,13]. These lizards have sex chromosomes of the XX/XY type, which are homologous to the sex chromosomes of most other pleurodont iguanians, and are involved in fusions in the species with low diploid numbers [12,14].

In this study we investigated the karyotype of *S. malachiticus,* which represents the clade with 2n=22. We obtained flow-sorted chromosome samples, which were used for FISH and low coverage sequencing to determine their homology with the chromosomes of *Anolis carolinensis,* a species which retains the ancestral karyotype of Iguania [15].

To check the hypothesis on the connection between chromosome fusion and evolution of recombination patterns, we applied an immunocytological approach to detect meiotic crossing over in synaptonemal complex (SC) spreads of *S. malachiticus* and *S. variabilis,* a related species with the unaltered ancestral karyotype (2n=34), and analyzed the recombination patterns using the *r* parameter [16]. This parameter is designed to measure recombination rate in meiosis by taking account of chromosome number, crossover number and crossover location. Its physical sense is the probability of two randomly selected loci to recombine in a meiotic act. It has two components: the interchromosomal, which reflects recombination via independent chromosome segregation, and the intrachromosomal, which reflects crossover recombination. We also studied sex chromosome synapsis and recombination to validate the FISH and sequencing data on their structure.

## Materials and methods

The male specimens of *S. malachiticus* and *S. variabilis* were obtained from the pet trade. To confirm the species identification, the 5’-fragment of the mitochondrial COI gene was sequenced using protocols and primers described previously [17].

The primary fibroblast cell cultures of *S. malachiticus* were obtained in the Laboratory of Animal Cytogenetics, the Institute of Molecular and Cellular Biology, Russia, using the protocols described previously [18,19]. All cell lines were deposited in the IMCB SB RAS cell bank (“The general collection of cell cultures”, 0310-2016-0002). Metaphase chromosome spreads were prepared from chromosome suspensions obtained from early passages of primary fibroblast cultures as described previously [20–22].

C-like DAPI staining was performed in the following way. The slides were incubated in 0.2 M HCl for 20 min at room temperature. Then they were kept in Ba(OH)_2_ solution at 55° C for 4 min, and then incubated in 2xSSC at 60° C for 60 min. Then they were washed in distilled water at room temperature, and stained with DAPI using the Vectashield mounting medium with DAPI (Vector Laboratories).

The SC spreads of *S. malachiticus* and *S. variabilis* were prepared and immunostained as described previously, using the antibodies to SYCP3, the protein of the lateral element of the SC; to MLH1, the mismatch-repair protein which marks mature recombination nodules; and to CENP, the centromere proteins [23].

The flow-sorted chromosome samples of *S. malachiticus* were obtained using the Mo-Flo^®^ (Beckman Coulter) high-speed cell sorter at the Cambridge Resource Centre for Comparative Genomics, Department of Veterinary Medicine, University of Cambridge, Cambridge, UK, as described previously [24]. The painting probes were generated from the DOP-PCR amplified samples by a secondary DOP-PCR incorporation of biotin-dUTP and digoxigenin-dUTP (Sigma) [25]. FISH was performed with standard techniques [26].

The preparations were analyzed with an Axioplan 2 Imaging microscope (Carl Zeiss) equipped with a CCD camera (CV M300, JAI), CHROMA filter sets, and the ISIS4 image processing package (MetaSystems GmbH). The brightness and contrast of all images were enhanced using Corel PaintShop Photo Pro X6 (Corel Corp).

For the preparation of the sorted chromosomes of *S. malachiticus* for sequencing, we used TruSeq Nano DNA Low Throughput Library Prep Kit (Illumina). Paired-end sequencing was performed on Illumina MiSeq using Reagent Kits v2, 600-cycles. The NGS data were deposited in NCBI SRA database (PRJNA616430). Sequencing data were processed using the “DOPseq_analyzer” pipeline [27, 28]. The following parameters were used: for read trimming “ampl” was set to “dop”, Illumina adapter trimming was enabled, additional cutadapt options “–trim-n –minimum-length 20” were specified. Reads were aligned to *A. carolinensis* genome (AnoCar2.0) downloaded from Ensembl (www.ensembl.org) using the BWA MEM algorithm with default parameters. Additional filters were minimum MAPQ = 20 and minimum alignment length = 20. For target region identification, scaffolds with chromosome assignments and scaffolds over 50 kb in size were used. The key parameter in the “DOPseq_analyzer” output to determine the target scaffolds and chromosomes is *pd_mean,* which is mean distance between the positions of the scaffold which are covered by the sequencing reads. The target scaffolds are characterized by lower *pd_mean* and contaminant scaffolds are characterized by higher *pd_mean.* The position of the scaffolds on the chromosomes of *A. carolinensis* was determined using the data on their synteny to the chicken chromosomes, obtained from UCSC Genome Browser (www.genome.ucsc.edu), and the previously obtained data on the synteny between the *Anolis* and chicken chromosomes [28,29].

## Results

### Species identification and the mitotic karyotype of S. malachiticus

The COI gene sequence confirmed that the specimens studied belong to *S. malachiticus* (GenBank MT140115) and *S. variabilis* (GenBank MT131783). The mitotic karyotype of *S. malachiticus* was typical for the 2n=22 clade of *Sceloporus* [12], and included six pairs of large metacentric chromosomes, medium-sized X and Y chromosomes, and four pairs of small metacentric chromosomes (Fig. 1a). The flow-sorted karyotype consisted of 12 peaks (Fig. 1b).

**Fig. 1.**
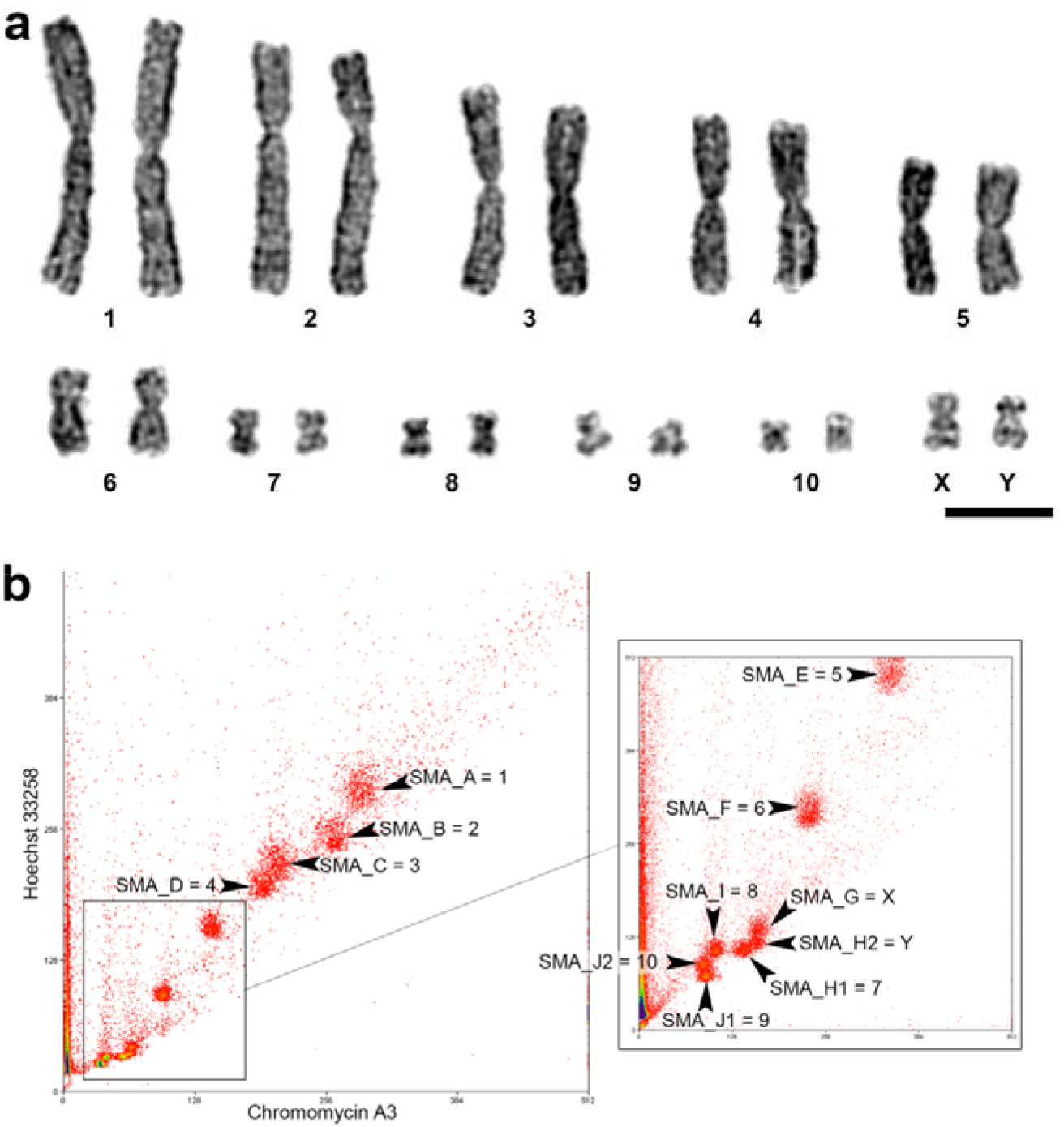
Karyotype of *S. malachiticus.* a: G-banded karyotype. Scale bar: 10 μm. b: flow-sorted karyotype. Peak IDs and contents are indicated by the arrowheads. X and Y axes: fluorescence intensity for each fluorochrome.

C-like DAPI staining showed DAPI-positive bands in the centromeres of each chromosome. The Y chromosome had more prominent band than the X chromosome. The chromosome 7 was totally heterochromatic (Fig. 2).

**Fig. 2.**
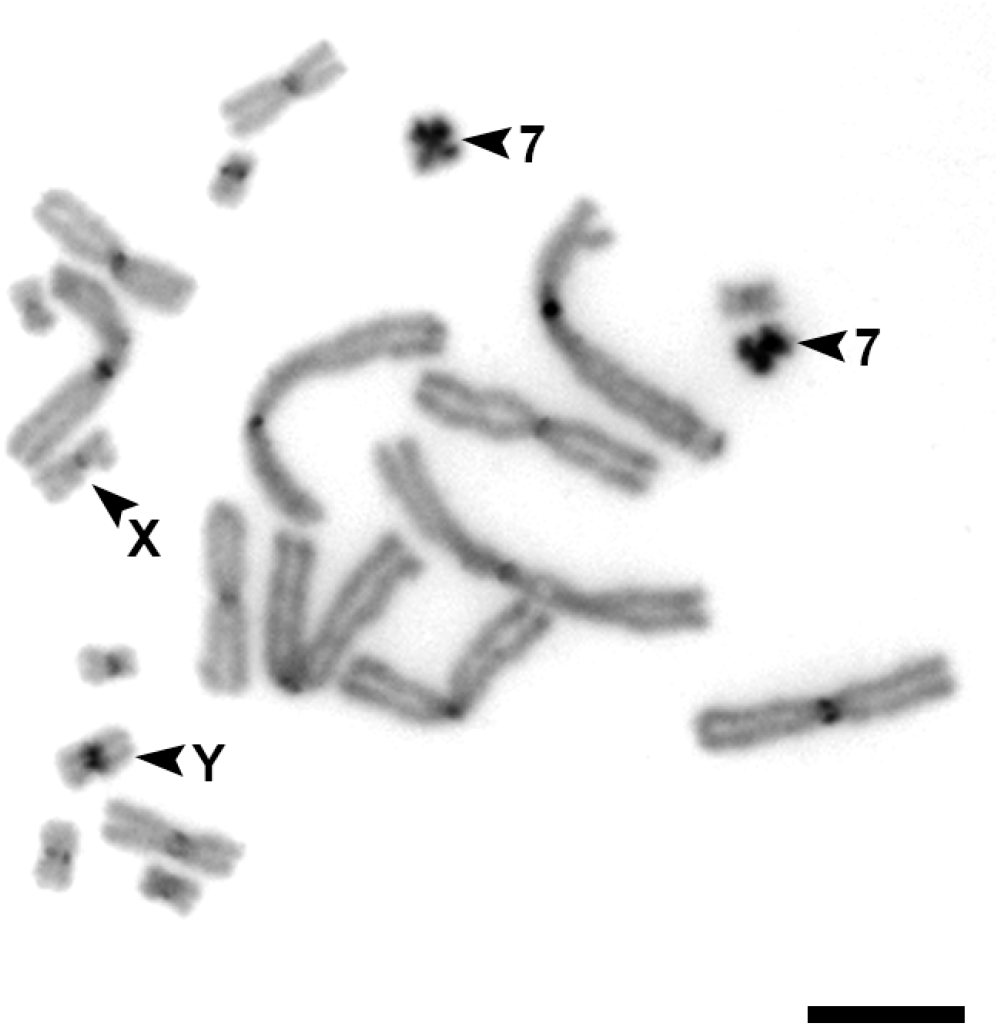
C-like DAPI staining of the *S. malachiticus* metaphase plate. Arrowheads indicate the heterochromatic chromosome 7 and the sex chromosomes. Scale bar: 10 μm.

### FISH with the flow-sorted chromosome-specific probes

FISH with the labelled probes derived from each peak was performed on the metaphase plates of the same specimen. The probes derived from most peaks hybridized with single chromosome pairs (Fig. 3). Two peaks, SMA_G and SMA_H2, both hybridized with the sex chromosomes, and were concluded to correspond to the chromosomes X and Y, respectively. The peak SMA_H2 (mostly consisting of the Y chromosome), in addition, showed weak hybridization with the similar-sized chromosome 7 (mostly contained in the peak SMA_H1) probably due to contamination during flow sorting (Fig. 3d).

**Fig. 3.**
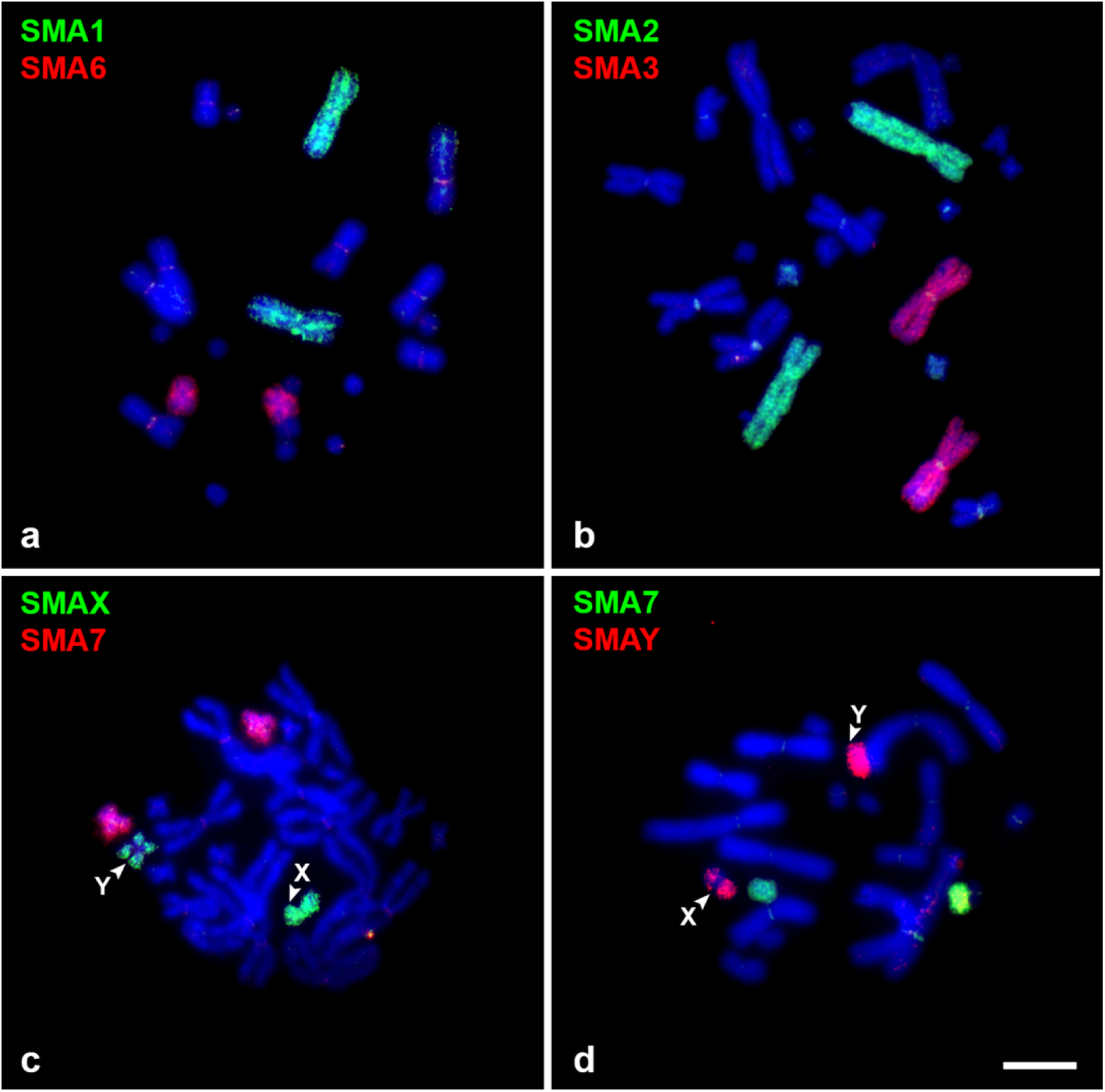
Examples of FISH with the flow-sorted chromosome-specific probes of the male *S. malachiticus* on its metaphase plates. Note that the sex chromosome-specific probes label the respective chromosomes across the whole length, but label the sex chromosomes of the opposite type (e.g., the SMA_X probe on the Y chromosome) only in the distal pseudoautosomal regions. Scale bar: 10 μm.

Both sex chromosome probes hybridized with the respective chromosomes along the whole length, but showed weaker or no hybridization in the medial, centromere-adjacent region of the homologous chromosome (Fig. 3 c, d). This pattern corresponds to the presence of two homologous pseudoautosomal regions in the distal parts of the sex chromosomes, and the differentiated non-recombining regions in the middle.

### Sequencing of the flow-sorted chromosome-specific probes

The NGS data analysis showed that the six largest macrochromosomes of *S. malachiticus* (SMA1-SMA6) correspond to the macrochromosomes of *A. carolinensis* (ACA1-ACA6), with two notable exceptions (Table 1, Supplementary file 1).

**Table 1.** The homology between the chromosomes of *S. malachiticus* and *A. carolinensis,* inferred from the NGS data.

| *S. malachiticus*<br>chromosomes | *A. carolinensis*<br>chromosomes |
| --- | --- |
| 1 | 1 |
| 2 | 2, 1 |
| 3 | 3 |
| 4 | 4 |
| 5 | 5 |
| 6 | 6, 1 |
| X | X, 11, 16, 17, 18 |
| Y | 11, 16, 17, 18 |
| 7 | 15 |
| 8 | 7, 10 |
| 9 | 9, 14 |
| 10 | 8, 12 |

In addition to high homology to ACA2, SMA2 shows homology with the segment located between the positions 204854915-205800612 of ACA1 (AnoCar2.0). In ACA1, this fragment represents an isolated segment of homology with the chicken chromosome 12 (GGA12), surrounded by the segments homologous to GGA3. It is notable that the segments homologous to GGA12 are mostly located on ACA2.

Similarly, SMA6, along with homology to ACA6, shows homology with the segment located between the positions 192604161-196661610 of ACA1 (AnoCar2.0). In ACA1, this genomic block is homologous to a segment of GGA2, and is also surrounded by the segments homologous to GGA3. The adjacent segments of GGA2 are homologous to ACA6.

The sex chromosomes of *S. malachiticus* contain the genomic blocks which are homologous to the chromosomes ACAX, ACA11, ACA16, ACA17, and ACA18. In the Y chromosome-specific DNA sample, 3 of 47 (6%) identified homologous ACA scaffolds belonged to ACAX, in contrast with 7 of 53 (13%) in the X chromosome. These scaffolds showed higher *pd_mean* than the scaffolds corresponding to other ACA chromosomes. They probably represent a contamination of the Y-sample by the X chromosome due to similar size.

The chromosomes SMA8, SMA9, and SMA10 correspond to the chromosomes ACA7+ACA10, ACA9+ACA14, and ACA8+ACA12, respectively. In the chromosome SMA7, only the genomic blocks homologous to ACA15 were found.

The data on the homology between the chromosomes of *S. malachiticus* and *A. carolinensis* are summarized in the Table 1 and Supplementary File 1.

### Synaptonemal complex analysis

The SC karyotype of *S. malachiticus* consisted of 11 metacentric bivalents, including the XY bivalent with misaligned centromeres and a lateral buckle on one of the homologues (Fig. 4a). This corresponds to the previously reported SC karyotype of a closely related species *S. undulatus* (Reed et al., 1990), and to the mitotic karyotype of *S. malachiticus* obtained in the present work. The recombination nodules, marked with the MLH1 protein, were located in the distal parts of the sex bivalent (Fig. 4b). The SC karyotype of *S. variabilis* consisted of 17 bivalents: 6 metacentric macrochromosomal bivalents and 11 microchromosomal bivalents (Fig. 4c). This corresponds to its previously known mitotic karyotype [30]. One of the microg-bivalents consisted of homologues of unequal lengths, with the larger homologue forming a lateral buckle (Fig. 4c,d). It was concluded to be the sex bivalent, basing on its similarity with the previously studied micro-sex bivalents in *Anolis* [31].

**Fig. 4.**
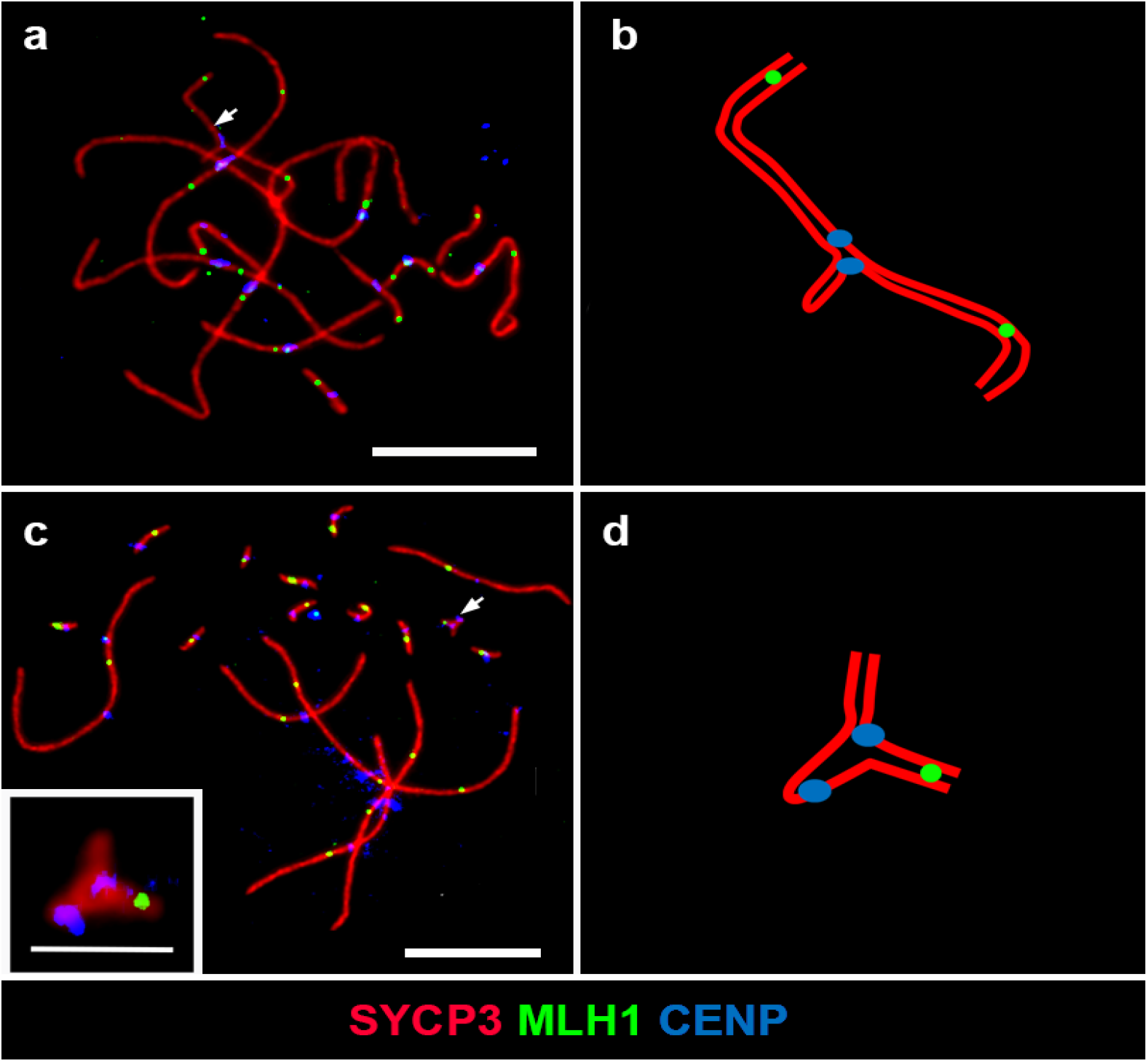
Synaptonemal complex analysis of *S. malachiticus* (a,b) and *S. variabilis* (c, d). a,c: immunofluorescent staining of SC spreads. The sex bivalents are indicated by arrows. Scale bars: 10 μm. Insert: the micro-sex bivalent of *S. variabilis*, scale bar: 2μm. b,d: schematic drawings of the sex bivalents.

The average crossover numbers were 15.6±2.3 per spread in *S. malachiticus* and 20.9±1.7 per spread in *S. variabilis*. However, the *r* parameter shows that the lower number of crossovers in *S. malachiticus* reflects only its lower chromosome number and not a change in crossing over patterns: the intra-chromosomal components of *r* are 0.032±0.005 in *S. malachiticus* and 0. 033±0.006 in *S. variabilis.* It is notable that both species have elevated crossover numbers near the centromeres of macro-autosomes, as shown previously by the analyses of meiotic metaphase chromosomes [12,14] (Fig. 5).

**Fig. 5.**
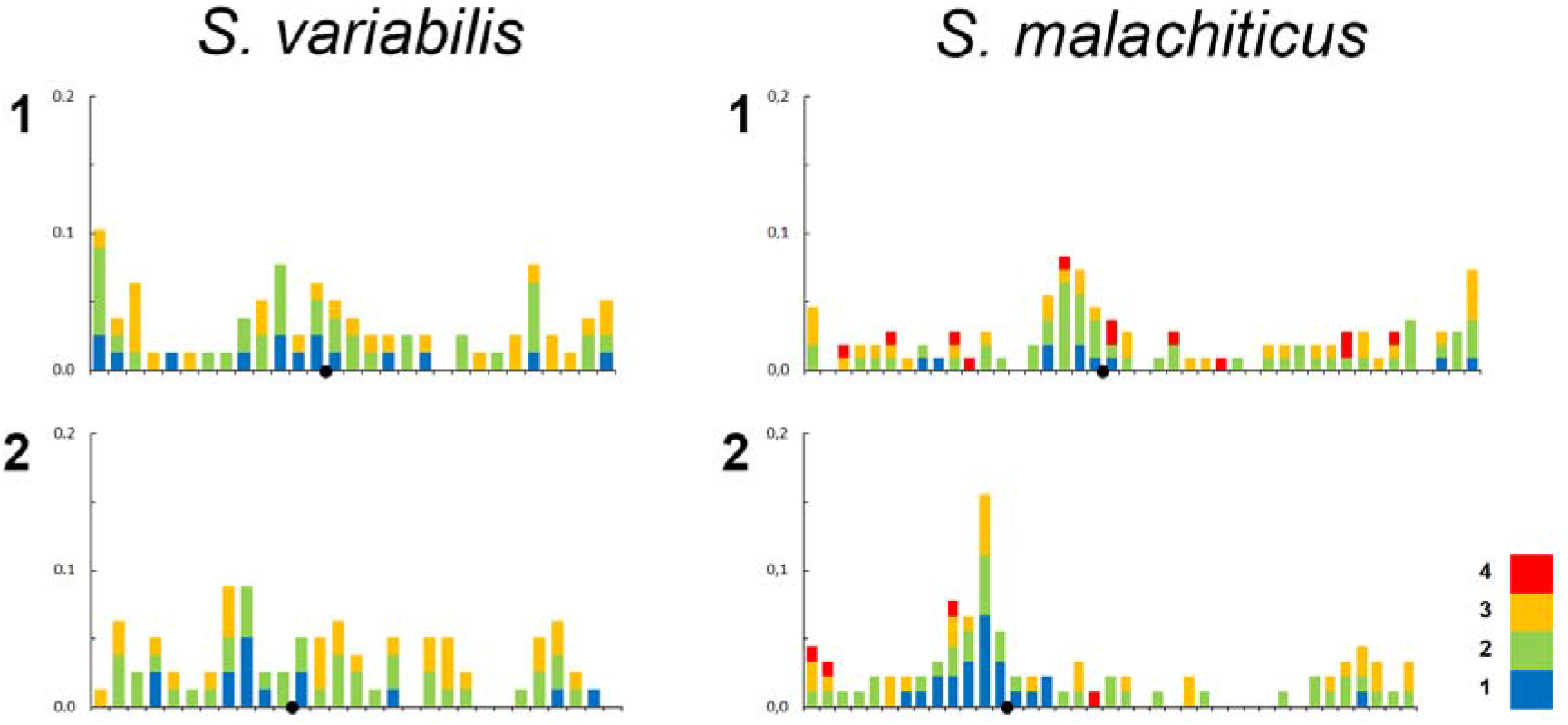
Numbers and distributions of the MLH1 foci on two largest chromosomes of *S. variabilis* and *S. malachiticus.* The x-axis shows the positions of MLH1 foci along the SCs in relation to the centromere (black circle). One scale division represents a segment of the average length of each SC equal to 1 μm. The y-axis shows the proportion of MLH1 foci in each interval. Different colors show bivalents with different MLH1 numbers, from 1 to 4.

## Discussion

The results of FISH and whole chromosome-specific DNA sequencing show that the macrochromosomes were stable during karyotypic evolution of *S. malachiticus,* in contrast to the microchromosomes. The segments of ACA1 which show homology to SMA2 and SMA6 are homologous to the segments of the chicken chromosomes GGA12 and GGA2. Most parts of these chicken chromosomes have homology with SMA2/ACA2 and SMA6/ACA6, respectively. This indicates that these segments of ACA1 most probably represent relatively recent translocations from proto-ACA2 and proto-ACA6, which occurred in the *Anolis* lineage and are not shared by *Sceloporus.* Alternatively, this might represent an assembly error in the *A. carolinensis* genome.

SMA7 is larger than other small autosomes, which contain homologues of two *A. carolinenesis* microchromosomes each (Fig. 1b, Table 1). However, its sequence shows homology only with ACA15. It is possible that SMA7 also contains genomic segments which are not covered or not assembled in the current version of the *A. carolinensis* genome, or is enlarged due to repeat accumulation. The latter explanation is supported by the high heterochromatinization of SMA7 (Fig. 2).

There are much data on the homology between the macrochromosomes of *A. carolinensis* and chromosomes of other squamates [6,7,32]. However, data for the microchromosomes are much less available. The established homologies between the microchromosomes of *A. carolinensis* and chromosomes of other squamates mainly concern the chromosomes which are involved in sex chromosome formation in different species.

The sex chromosome-autosome fusions can become fixed in different ways. According to the most widespread model of sex chromosomes evolution, the translocations of loci with sexual antagonistic alleles on the sex chromosomes are positively selected, and such loci accumulate in the sex chromosomes [33,34]. Another model shows that such fusions could be favored in cases of selection for heterozygosity [35]. There is also a model which implies that the fusions are deleterious and are fixed by drift [36]. Thus, the predisposition of fusions between certain autosomes and sex chromosomes, irrespectively of the genomic origin of the sex chromosomes, to become fixed, may arise if sex linkage or high heterozygosity are particularly beneficial for certain loci, or if these fusions are the least deleterious.

The sex chromosomes in *S. malachiticus* experienced more fusions than the autosomes. Both FISH and crossover mapping data show the presence of two distal pseudoautosomal regions, and a lack of homology between the X and Y chromosomes in the median parts. This indicates that the autosomal fragments translocated onto the sex chromosomes from both sides. However, our methods do not allow determination of the relative positions of each syntenic block inside the sex chromosomes.

Of the fusions which we identified in the sex chromosomes of *S. malachiticus,* several appear to be homoplasies, as they appear independently in different squamate lineages. Namely, the combination of the homologues of ACAX+ACA11 occur independently in *Ctenonotus* (Dactyloidae, Pleurodonta) [37,38] and in the Z chromosome of *Paroedura* (Gekkonidae) [39]; the combination of the homologues of ACAX+ACA18 occur in *Norops* (Dactyloidae, Gekkota) [28]; and the combination of the homologues of ACA11+ACA16 occur in the Z chromosome of Lacertidae [40]. It is also notable that both in *Norops* and *Sceloporus* the sex chromosomes stand out by undergoing more fusions than autosomes. However, the homologues of ACA9 and ACA12, which constitute the sex chromosomes of *Norops* along with ACAX and ACA18, are not fused with the sex chromosomes and with each other in *S. malachiticus*. The fusion of the homologues of ACA15 and ACA16, which created the chromosome 12 of *Norops* [28], did not take place in *S. malachiticus.*

Thus, although we detected some repeated microchromosome fusions, the paucity of the available comparative data prevents us from determining whether certain combinations of microchromosomes fuse significantly more frequently than others. The repeated involvement of the homologues of ACA11 in the formation of sex chromosomes requires special attention, since this chromosome is homologous to the sex chromosome system of therian mammals. More comprehensive understanding of our data and more detailed comparative analysis will be available with the emergence of more syntenic maps of squamates with fused microchromosomes. In some of them, the existing syntenic maps cover only the homologues of the *A. carolinensis* macrochromosomes, as in lacertids [6] and geckos [7] or have too small numbers of markers, as in tuatara [41]. For others, *i.e.* skinks, syntenic maps do not exist at all [42].

We did not confirm the hypothesis that *S. malachiticus* could have lower crossing over rates than the related bimodal species. On the contrary, both species of *Sceloporus* show similar intra-chromosomal components of r, which are higher than in two previously studied bimodal *Anolis* species [23]. Thus, at least in the studied species of Pleurodonta, the crossover patterns depend on the phylogenetic position of the species, and not on the structure of its karyotype. The high *r* values in *Sceloporus* are due to the centromeric peaks of crossover distribution, in contrast with *Anolis*, which demonstrate mostly distally located crossovers. As shown by Veller et al., [16], median crossovers contribute much more to the effective recombination rate than the distal ones. It is highly unusual for vertebrates to have crossover peaks near the centromeres: the more common pattern is a reduction of crossover rate in the centromeric regions, which is called the “centromere effect” [43]. The physiological mechanisms and possible biological significance of the altered crossover patterns in *Sceloporus* deserve further study.

## Acknowledgements

We thank the Microscopic Center of the Siberian Branch of the Russian Academy of Sciences for granting access to microscopic equipment. We thank I. Kichigin for help in NGS data analysis and K. Petrova for assisting in G-banding of *S. malachiticus* chromosomes. This work was supported by the research grant #19-54-26017 from the Russian Foundation for Basic Research, the research grant #19-14-00050 from the Russian Science Foundation, the research grant #AAAA-A 17-117071240065-4 from the Ministry of Science and Higher Education (Russia) via the Institute of Cytology and Genetics.

## References

1. Morescalchi A. 1977 Phylogenetic Aspects of Karyological Evidence. In Major Patterns in Vertebrate Evolution, pp. 149–167. Springer US. (doi:10.1007/978-1-4684-8851-7_7)

2. Olmo E, Signorino G. 2005 Chromorep: a reptiles chromosomes database. www.chromorep.univpm.it

3. Morescalchi A. 1980 Evolution and karyology of the amphibians. Bolletino di Zool. 47, 113–126. (doi:10.1080/11250008009438709)

4. Braasch I et al. 2016 The spotted gar genome illuminates vertebrate evolution and facilitates human-teleost comparisons. Nat. Genet. Vol. 48, 427–437. (doi:10.1038/ng.3526)

5. Uno Y, Nishida C, Tarui H, Ishishita S, Takagi C. 2012 Inference of the Protokaryotypes of Amniotes and Tetrapods and the Evolutionary Processes of Microchromosomes from Comparative Gene Mapping. PLoS One 7. (doi:10.1371/journal.pone.0053027)

6. Srikulnath K, Matsubara K, Uno Y, Nishida C, Olsson M, Matsuda Y. 2014 Identification of the linkage group of the Z sex chromosomes of the sand lizard (Lacerta agilis, Lacertidae) and elucidation of karyotype evolution in lacertid lizards. Chromosoma 123, 563–575. (doi:10.1007/s00412-014-0467-8)

7. Srikulnath K, Uno Y, Nishida C, Ota H, Matsuda Y. 2015 Karyotype reorganization in the Hokou Gecko (Gekko hokouensis, Gekkonidae): The process of microchromosome disappearance in Gekkota. PLoS One 10. (doi:10.1371/journal.pone.0134829)

8. Morescalchi A. 1977 Adaptation and Karyotype in Amphibia. Ital. J. Zool. 44, 287–294. (doi:10.1080/11250007709430182)

9. Charlesworth D. 2016 The status of supergenes in the 21st century: recombination suppression in Batesian mimicry and sex chromosomes and other complex adaptations. Evol. Appl. 9, 74–90. (doi:10.1111/eva.12291)

10. Deakin JE, Ezaz T. 2019 Understanding the Evolution of Reptile Chromosomes through Applications of Combined Cytogenetics and Genomics Approaches. Cytogenet Genome Res 157, 7–20. (doi:10.1159/000495974)

11. Sigeman H, Ponnikas S, Chauhan P, Dierickx E, De Brooke M, Hansson B. 2019 Repeated sex chromosome evolution in vertebrates supported by expanded avian sex chromosomes. Proc. R. Soc. B Biol. Sci. 286, 20192051. (doi:10.1098/rspb.2019.2051)

12. Hall WP. 2009 Chromosome variation, genomics, speciation and evolution in Sceloporus lizards. Cytogenet. Genome Res. 127, 143–165. (doi:10.1159/000304050)

13. Leaché AD, Banbury BL, Linkem CW, Nieto-Montes De Oca A. 2016 Phylogenomics of a rapid radiation: is chromosomal evolution linked to increased diversification in north american spiny lizards (Genus Sceloporus)? BMC Evol. Biol. 16, 63. (doi:10.1186/s12862-016-0628-x)

14. Reed KM, Sudman PD, Sites JW, Greenbaum IF. 1990 Synaptonemal Complex Analysis of Sex Chromosomes in Two Species of Sceloporus. Copeia, 1122–1129. (doi:10.2307/1446497)

15. Deakin JE et al. 2016 Anchoring genome sequence to chromosomes of the central bearded dragon (Pogona vitticeps) enables reconstruction of ancestral squamate macrochromosomes and identifies sequence content of the Z chromosome. BMC Genomics 17, 447. (doi:10.1186/s12864-016-2774-3)

16. Veller C, Kleckner N, Nowak MA. 2019 A rigorous measure of genome-wide genetic shuffling that takes into account crossover positions and Mendel’s second law. Proc. Natl. Acad. Sci. U. S. A. 116, 1659–1668. (doi:10.1073/pnas.1817482116)

17. Nagy ZT, Sonet G, Glaw F, Vences M. 2012 First large-scale DNA barcoding assessment of reptiles in the biodiversity hotspot of madagascar, based on newly designed COI primers. PLoS One 7. (doi:10.1371/journal.pone.0034506)

18. Stanyon R, Galleni L. 1991 Italian Journal of Zoology A rapid fibroblast culture technique for high resolution karyotypes A rapid fibroblast culture technique for high resolution karyotypes. Ital. J. Zool. 58, 81–83. (doi:10.1080/11250009109355732)

19. Romanenko SA et al. 2015 Segmental paleotetraploidy revealed in sterlet (Acipenser ruthenus) genome by chromosome painting. Mol. Cytogenet. 8, 90. (doi:10.1186/s13039-015-0194-8)

20. Yang F, O’brien PCM, Milne BS, Graphodatsky AS, Solanky N, Trifonov V, Rens W, Sargan D, Ferguson-Smith MA. 1999 A Complete Comparative Chromosome Map for the Dog, Red Fox, and Human and Its Integration with Canine Genetic Maps. Genomics 62, 189–202. (doi:10.1006/geno.1999.5989)

21. Graphodatsky A et al. 2000 Comparative cytogenetics of hamsters of the genus Calomyscus. Cytogenet. Cell Genet:. 88, 296–304. (doi:10.1159/000015513)

22. Graphodatsky AS, Yang F, O’brien PCM, Perelman P. 2001 Phylogenetic implications of the 38 putative ancestral chromosome segments for four canid species Create new project ‘Ultrasonic energy transport proved using femtosecond laser pulses’ View project Cytogenomics in Amazonian bats View project. Cytogenet. Genome Res. 92, 243–247. (doi:10.1159/000056911)

23. Lisachov AP, Tishakova K V., Tsepilov YA, Borodin PM. 2019 Male Meiotic Recombination in the Steppe Agama, Trapelus sanguinolentus (Agamidae, Iguania, Reptilia). Cytogenet. Genome Res. 157, 107–114. (doi:10.1159/000496078)

24. Yang F, Carter NP, Shiu L, Ferguson-Smith MA. 1995 A comparative study of karyotypes of muntjacs by chromosome painting. Chromosoma 103, 642–652. (doi:10.1007/BF00357691)

25. Telenius H, Carter NP, Bebb CE, Nordenskjöld M, Ponder BAJ, Tunnacliffe A. 1992 Degenerate oligonucleotide-primed PCR: General amplification of target DNA by a single degenerate primer. Genomics 13, 718–725. (doi:10.1016/0888-7543(92)90147-K)

26. Liehr T, Kreskowski K, Ziegler M, Piaszinski K, Rittscher K. 2016 The Standard FISH Procedure. In Springer Protocols Handbooks. pp. 109–118. Springer Berlin Heidelberg. (doi:10.1007/978-3-662-52959-1_9)

27. Makunin AI et al. 2016 Contrasting origin of B chromosomes in two cervids (Siberian roe deer and grey brocket deer) unravelled by chromosome-specific DNA sequencing. BMC Genomics 17, 618. (doi:10.1186/s12864-016-2933-6)

28. Kichigin IG et al. 2016 Evolutionary dynamics of Anolis sex chromosomes revealed by sequencing of flow sorting-derived microchromosome-specific DNA. Mol. Genet. Genomics 291, 1955–1966. (doi:10.1007/s00438-016-1230-z)

29. Alföldi J et al. 2011 The genome of the green anole lizard and a comparative analysis with birds and mammals. Nature 477, 587–591. (doi:10.1038/nature10390)

30. Porter CA, Haiduk MW, Queiroz K DE. 1994 Evolution and Phylogenetic Significance of Ribosomal Gene Location in Chromosomes of Squamate Reptiles. Copeia, 302–313. (doi:10.2307/1446980)

31. Lisachov AP, Trifonov VA, Giovannotti M, Ferguson-Smith MA, Borodin PM. 2017 Immunocytological analysis of meiotic recombination in two anole lizards (Squamata, Dactyloidae). Comp. Cytogenet. 11, 129–141. (doi:10.3897/CompCytogen.v11i1.10916)

32. Srikulnath K, Uno Y, Nishida C, Matsuda Y. 2013 Karyotype evolution in monitor lizards: Cross-species chromosome mapping of cDNA reveals highly conserved synteny and gene order in the Toxicofera clade. Chromosom. Res. 21, 805–819. (doi:10.1007/s10577-013-9398-0)

33. Rice WR. 1987 The Accumulation of sexually Antagonistic genes as a selective agent promoting the evolution of reduced recombination between primitive sex chromosomes. Evolution (N. Y). 41, 911–914. (doi:10.1111/j.1558-5646.1987.tb05864.x)

34. Charlesworth D, Charlesworth B. 1980 Sex differences in fitness and selection for centric fusions between sex-chromosomes and autosomes. Genet. Res. 35, 205–214. (doi:10.1017/S0016672300014051)

35. Charlesworth B, Wall JD. 1999 Inbreeding, heterozygote advantage and the evolution of neo-X and neo-Y sex chromosomes. Proc. R. Soc. London. Ser. B Biol. Sci. 266, 51–56. (doi:10.1098/rspb.1999.0603)

36. Pennell MW, Kirkpatrick M, Otto SP, Vamosi JC, Peichel CL, Valenzuela N, Kitano J. 2015 Y Fuse? Sex Chromosome Fusions in Fishes and Reptiles. PLoS Genet. 11. (doi:10.1371/journal.pgen.1005237)

37. Giovannotti M et al. 2017 New insights into sex chromosome evolution in anole lizards (Reptilia, Dactyloidae). Chromosoma 126, 245–260. (doi:10.1007/s00412-016-0585-6)

38. Lisachov AP, Makunin AI, Giovannotti M, Pereira JC, Druzhkova AS, Caputo Barucchi V, Ferguson-Smith MA, Trifonov VA. 2019 Genetic Content of the Neo-Sex Chromosomes in Ctenonotus and Norops (Squamata, Dactyloidae) and Degeneration of the Y Chromosome as Revealed by High-Throughput Sequencing of Individual Chromosomes. Cytogenet. Genome Res. 157, 115–122. (doi:10.1159/000497091)

39. Rovatsos M, Farkacová K, Altmanová M, Johnson Pokorná M, Kratochvíl L. 2019 The rise and fall of differentiated sex chromosomes in geckos. Mol. Ecol. 28, 3042–3052. (doi:10.1111/mec.15126)

40. Rovatsos M, Vukić J, Kratochvíl L. 2016 Mammalian X homolog acts as sex chromosome in lacertid lizards. Heredity 117(1), 8–13. (doi:10.1038/hdy.2016.18)

41. O’Meally D, Miller H, Patel HR, Marshall Graves JA, Ezaz T. 2010 The first cytogenetic map of the tuatara, sphenodon punctatus. Cytogenet. Genome Res. 127, 213–223. (doi:10.1159/000300099)

42. Lisachov AP, Poyarkov N, Pawangkhanant P, Borodin P, Srikulnath K. 2018 New karyotype of Lygosoma bowringii (Günther, 1864) suggests cryptic diversity. Herpetol. Notes 11, 1083–1088.

43. Nambiar M, Smith GR. 2016 Repression of harmful meiotic recombination in centromeric regions. Semin Cell Dev Biol 54, 188–197. (doi:10.1016/j.semcdb.2016.01.042)

